# Satellite cell-derived extracellular vesicles reverse peroxide-induced mitochondrial dysfunction in myotubes

**DOI:** 10.1101/2020.03.04.977280

**Authors:** Kyle T. Shuler, Brittany E. Wilson, Eric R. Muñoz, Andrew D. Mitchell, Joshua T. Selsby, Matthew B. Hudson

## Abstract

Satellite cells (SCs) are muscle-specific stem cells that have a central role in muscle remodeling. Despite their therapeutic potential, SC-based therapies have been met with numerous logistical challenges, limiting their ability to effectively treat systemic muscle diseases, such as Duchenne muscular dystrophy (DMD). Delivery of SC-derived extracellular vesicles (SC-EVs) may unlock the potential offered by SCs and overcome their numerous limitations.

**Purpose:** The purpose of this investigation was to determine the extent to which SC-EVs could restore mitochondrial function in cultured myotubes following oxidative injury.

**Methods:** SC-EVs were isolated from cultured SCs from C57 mice and quantified using nanoparticle tracking analysis (NTA). C2C12 myotubes were cultured and divided into four treatment groups: untreated control, treated for 24 h with SC-EV, 24 h exposure to 50 μM H_2_O_2_ followed by a 24 h recovery period with no treatment, or 24 h exposure to 50 μM H_2_O_2_ followed by a 24 h treatment with SC-EV. Inter-group differences in mitochondrial function were assessed via one-way ANOVA with Tukey post hoc analysis (p<0.05).

**Results:** Given the seeding density used, we calculated that each SC releases approximately 2.35 × 10^5^ ± 3.10 × 10^4^ EVs per 24 h. Further, using fluorescent microscopy, we verified SC-EVs deliver cargo into myotubes, some of which was localized to the mitochondria. H_2_O_2_ exposure resulted in a 42% decline in peak mitochondrial respiration (p=0.0243) as well as a 46% reduction in spare respiratory capacity (p=0.0185) relative to the untreated control group. Subsequent treatment with SC-EVs (3.12×10^8^ SC-EV; 24 h) following H_2_O_2_ exposure restored 76% of peak mitochondrial respiration (p=0.0187) and 84% of spare respiratory capacity in the damaged myotubes (p=0.0198). SC-EVs did not affect mitochondrial function in the undamaged myotubes.

**Conclusion:** Collectively, these data demonstrate SC-EVs may represent a novel therapeutic approach for treatment of myopathies associated with mitochondrial dysfunction.

## Introduction

Satellite cells (SCs) are skeletal muscle stem cells and play a central role in skeletal muscle remodeling [1]. Upon injury to muscle, SCs are activated, proliferate, and differentiate into myoblasts and then fuse into the preexisting myotubes to facilitate regeneration. Due to their role in regeneration and repair, SCs may provide therapeutic benefit to a range of muscle disorders and pathologies [2]. Although current strategies in SC-based therapies have focused primarily on the isolation, culture, and transplantation of SCs in the treatment of muscular dystrophies, these cells may also hold therapeutic potential in other conditions, such as cachexia, sarcopenia, disuse muscular atrophy, and muscle trauma [3–7]. Although initially promising, there are limitations to this strategy [8, 9]. For example, since skeletal muscle is the most abundant tissue in humans, a very high number of SCs are likely required to treat systemic muscle conditions, though the number of SCs required to achieve a lasting therapeutic benefit is equivocal [2]. Further, generating the large number of SCs needed for therapeutics requires expanding SCs *ex vivo*; however, the capacity to expand isolated SCs is limited before myogenic potential rapidly declines [10].

Extracellular vesicles (EVs) are small, lipid-coated vesicles that contain molecular cargo and are generally classified into overlapping size ranges as exosomes (30-150nm), microvesicles (100-800nm), and large EVs (apoptotic bodies, 200nm-5µm), with their more distinct characteristics being differences in biogenesis and molecular contents. EVs participate in paracrine and endocrine-like signaling via the selective packaging of specific molecular cargo that are delivered into nearby and distant recipient cells. [11]. Most, if not all cell types, including muscle cells, release EVs [12, 13]. Recent evidence indicates EVs play critical roles in allowing SCs to respond to stimuli as well as interact with other cell types within the muscle niche [14–16]. Additionally, EVs released by cardiosphere-derived cells transiently restore dystrophin in the myocardium and skeletal muscle of dystrophic mice, potentially via delivery of miRNA 148a [17].

Feasibly, satellite cell-derived extracellular vesicles (SC-EVs) may provide therapeutic benefits for myopathies and may help overcome many of the limitations of traditional SC-mediated therapeutic approaches; however, limited information exists regarding SC-EVs. Therefore, the purpose of this investigation was to characterize SC-EVs and examine potential therapeutic benefits *in vitro.* Importantly, mitochondrial damage has been causatively linked to impaired SC function [18, 19] and impaired mitochondrial function is known to play a role in muscle atrophy, and/or weakness occurring in muscular dystrophy [20–22], cancer cachexia [23], disuse atrophy [24, 25], mechanical ventilation-induced diaphragm weakness [26–30], sarcopenia [31, 32], and chronic kidney disease [33]. With the established links between SC function and mitochondrial function [34], we also investigated the extent to which SC-EVs could attenuate mitochondrial dysfunction. We hypothesized that SCs release high numbers of EVs that rapidly enter myotubes and attenuate oxidative stress-induced mitochondrial dysfunction.

## Results

### Characterization of SC-EVs

To determine the size and number of extracellular vesicles (EVs) released from satellite cells (SCs), the media was collected from cultured SCs 24 h after plating the cells (1.0-2.0 × 10^4^ cells/cm^2^). EVs were then isolated from media and quantified via nanoparticle tracking analysis (NTA). Over this 24 h period each SC released 2.35 × 10^5^ ± 3.10 × 10^4^ EVs, calculated using the data derived from NTA and the seeding density of the SCs. Since numerous EV subpopulations exist, we examined the size distribution of the satellite cell-derived extracellular vesicles (SC-EVs). We discovered SC-EVs have a mean size of 125.7 ± 1.7 nm and modal size of 99.30 ± 2.6 nm, indicating SCs primarily release EVs in the size range of exosomes and microvesicles (Figure 1).

### SC-EVs rapidly deliver protein into myotubes

After establishing that SCs release a substantial amount of EVs, we worked to demonstrate that these EVs could deliver proteins to recipient cells. Importantly, for these experiments we used differentiated myotubes as recipient cells to better model adult muscle fibers, *in vivo*. SC-EVs were incubated with carboxyfluorescein succinimidyl ester (CFSE) to label luminal proteins and were then delivered to myotubes by delivering the labeled EVs directly to the culture media. Following a 24 h incubation, myotubes were visually inspected using fluorescent microscopy. There appeared to be an abundance of fluorescent puncta within the cultured myotubes, suggesting the cells were able to uptake the CFSE-labeled protein from the SC-EVs (Figure 2 b). Using the same technique previously described, we next quantified the kinetics of SC-EV protein delivery by incubating SC-EVs in the culture media of myotubes for various durations ranging from 5 min to 48 h.

Fluorescent signal from the CFSE dye was then quantified to measure the amount of protein uptake in the myotubes. Remarkably, significant delivery of SC-EV protein was detected within the myotubes as early as 2 h after exposure (p= 0.0141; Figure 2 h) and for upwards of 48 h after exposure (p=0.0071; Figure 2 j & 3), indicating that the cargo was not rapidly degraded once delivered into the myotubes. The greatest amount of labeled protein was observed at 24 h after exposure (p=0.0004; Figure 2 k). All groups were compared to untreated control myotubes.

### SC-EV protein colocalizes to mitochondria in myotubes

Once we established SC-EV protein uptake was greatest at 24 h, we next examined the localization of the protein inside the myotubes. CFSE-labeled SC-EVs were incubated in the culture media of myotubes for 24 h. The myotubes were then stained with a mitochondria-specific dye. Fluorescent microscopy revealed what appeared to be overlapping signal for a portion of the labeled SC-EV protein and mitochondria (yellow) as well as fluorescent puncta juxtaposed to portions of the mitochondrial network (Figure 2 c). This suggested that the SC-EV protein cargo may interact with portions of the mitochondrial network.

### SC-EVs reverse peroxide-induced mitochondrial dysfunction

As it was clear that SC-EVs effectively delivered protein to recipient muscle cells, we next explored the potential of these EVs to attenuate dysfunction caused by acute oxidative stress. Mitochondrial respiration was measured in C2C12 myotubes under control conditions and following a 24 h treatment with 50µM hydrogen peroxide (H_2_O_2_). H_2_O_2_ exposure resulted in a 42% decline in peak mitochondrial respiration (p=0.0243; Figure 4 a & d) as well as a 46% reduction in spare respiratory capacity relative to the untreated control group (p=0.0185; Figure 4 a & e). Subsequent treatment with SC-EVs (3.12×10^8^ SC-EV; 24 h) following H_2_O_2_ exposure resulted in a 76% increase in peak mitochondrial respiration (p=0.0187; Figure 4 a & d) and 84% increase in spare respiratory capacity in the damaged myotubes (p=0.0198; Figure 4 a & e), reversing H_2_O_2_- induced dysfunction. We also included myotubes treated with SC-EVs without oxidative insult. Peak mitochondrial respiration and spare respiratory capacity were similar to control myotubes (Figure 4 d & e). Further, the ratio of oxygen consumption rate to extracellular acidification rate indicates that treatment with SC-EVs restored the energetic phenotype of the damaged myotubes to that of the control group (Figure 4 b). There was no significant differences between the basal respiration values of each group (Figure 4 c).

## Discussion

Use of satellite cells (SCs) as therapeutic agents has been widely investigated [2, 6, 35]. The therapeutic application of these cells is attributed to their ability to differentiate into myoblasts and fuse into damaged fibers to support regeneration [5]. Evidence suggests extracellular vesicles (EVs) released from mesenchymal stem cells confer many of the same regenerative benefits as the cells themselves [36]. Given the potential of satellite cell-derived extracellular vesicles (SC-EVs) to impact adult fibers, the purpose of this investigation was to describe and quantify the complement of SC-EVs and determine the extent to which SC-EVs attenuate mitochondrial dysfunction caused by acute oxidative stress. We found SCs release approximately 235,000 EVs/day and that adult myotubes take up large amounts of SC-EVs. Finally, we also discovered EVs reverse mitochondrial dysfunction caused by oxidative stress.

Mitochondria are critical in cellular energy production as well as the regulation of overall cellular viability [37]. Mitochondrial dysfunction plays a central role in numerous muscular disorders and pathologies, including DMD [38, 39]. For example, work by Hughes et al (2019) demonstrated that mitochondrial dysfunction, attributed to impaired oxidative phosphorylation and elevated oxidant production, contributes to early stage pathology in the D2.*mdx* mouse model [40]. Additionally, in the mdx mouse model, mitochondrial dysfunction is one of the earliest cellular consequences of DMD and is associated with the impaired ability of dystrophic muscle cells to respond to sarcolemmal damage [39]. Adverse alterations in mitochondrial morphology and function are also known to play a central role in the pathology of muscle dysfunction and wasting in murine models of cancer cachexia, attributed to decreased ATP production and uncoupling of oxidative phosphorylation [41]. It is also well-documented that mitochondrial dysfunction contributes to disuse atrophy [24]. For example, work by Powers et al (2011) demonstrates that a mitochondrial-targeted antioxidant was able to ameliorate ventilation-induced diaphragm weakness [25]. Given that mitochondrial dysfunction is associated with or contributes to an array of muscle pathologies, SC-EVs may alleviate a significant disease burden and provide therapeutic relief to a large number of patients. This approach has distinct advantages over cell transplantation including, most notably, the abundance of EVs released and the efficiency with which they deliver molecular cargo [42] [43].

While it is clear that SC-EVs attenuate mitochondrial dysfunction caused by acute oxidative stress, the underlying mechanism is decidedly less apparent. Speculatively, these EVs may deliver some factor (RNA and/or protein) that serves to reverse H_2_O_2_-mediated mitochondrial damage and restore mitochondrial function. At present, the identity of this factor or factors remains unknown. Given the 24 h period of SC-EV exposure to myotubes there is ostensibly sufficient duration to allow translation of delivered RNA or incorporation of delivered protein into the appropriate cellular compartments, including the mitochondria, themselves. Gartz et al (2019) demonstrated that exosomes from wild type and dystrophin-deficient induced pluripotent stem cell-derived cardiomyocytes exert cardioprotective effects on dystrophic cardiomyocytes [44]. The authors attributed these findings to a decrease in reactive oxygen species and delay in the formation of the mitochondrial transition pore following oxidative injury with H_2_O_2_, mediated by ERK1/2 and p38 MAPK signaling. Alternatively, SC-EVs may deliver intact mitochondria to recipient cells. This mechanism is attractive as EV-mediated delivery of mitochondria to damaged neurons has been demonstrated previously [45]. Our data indicated that SC-EVs delivered to undamaged myotubes did not alter mitochondrial respiration. If mitochondria were being delivered, we would expect to see increased respiration in this group. Given that there was only an increase in mitochondrial respiration in the group that was damaged first, it appears that the SC-EVs are delivering cargo that helps restore mitochondrial function but does not expand the overall mitochondrial capacity of otherwise healthy myotubes.

In summary, these data suggest that SCs release large amounts of EVs of multiple sizes, which are able to deliver molecular cargo to mature myotubes. We also discovered that these SC-EVs reverse oxidative stress-mediated mitochondrial dysfunction and restore the energetic phenotype of the damaged myotubes. Given this profound therapeutic response and the role of oxidative stress in a host of muscle pathologies, SC-EVs may be an effective, simple, safe, and scalable therapeutic approach. Future studies are needed to investigate the physiological roles of SC-EVs in the treatment of these pathologies as well as an identification of the mechanisms that underlie SC-EV-mediated restoration of mitochondrial function.

## Material and Methods

### Animals

All animal procedures were approved by the IACUC at the University of Delaware. C57BL/6 mice (n=8, 4-6 weeks of age, mixed male and female) were anesthetized using isoflurane gas throughout the dissection protocol.

### Isolation of Satellite Cells

All the muscles of the lower limb were removed from both limbs and stored on ice in wash buffer, consisting of 5% horse serum and 1% penicillin-streptomycin in DMEM. The muscles were then minced into a slurry, containing pieces approximately 3 mm in diameter [46]. The tissue was then transferred to dissociation buffer (3.3 mg/ml collagenase II in wash buffer) and incubated at 37°C for 1 h, while being mixed every 5-10 min. This was followed by a wash step and centrifugation at 500 x g for 5 min before removing the dissociation buffer and adding stock collagenase II (4.3 mg/ml) (Worthington, Lakewood, USA) and stock dispase (6.0 mg/ml) (Gibco, Gaithersburg, USA) to the solution. This solution was then incubated for an additional 30 min at 37°C while mixing. A 10 ml, 20 gauge needle was then used to triturate the solution. This was followed by an additional wash before the solution was run through a 40 µm filter. Two washes through the filter were then performed. The cells were then pelleted by centrifugation at 500 x g for 5 min at 22°C and all of the media was aspirated off. A Satellite Cell MACS cocktail (Satellite Cell Isolation Kit, Bergisch Gladbach, Germany) was then used to isolate the SCs from the cell suspension via magnetic separation.

### Satellite Cell Culture

Following isolation, the SCs were seeded on Matrigel-coated (Corning, Corning, USA) 6-well plates at a seeding density of 1.5×10^4^-2×10^4^ cells/cm^2^. The cells were grown in vesicle-free expansion media, consisting of 20% vesicle-free fetal bovine serum, 1% penicillin-streptomycin, 5 ng/ml basic-fibroblast growth factor (Progen, Heidleberg, Germany), and equal parts DMEM and Ham’s F10 mix. The SCs were cultured for six days in a 37°C incubator with 5% CO_2_, with media being replaced and collected each day.

### C2C12 Culture

C2C12 myoblasts were cultured in various vessels depending on downstream application. The cells were seeded at density of 1.0 × 10^4^ cells/cm^2^ and were then grown in a 37°C incubator with 5% CO_2_. The growth media used contained 10% fetal bovine serum and 1% penicillin-streptomycin in DMEM. The cells were proliferated until they were 90-100% confluent before being switched to differentiation media containing 2% horse serum and 1% p/s in DMEM. Following 5 d in differentiation media, the myoblasts had adequately fused to form myotubes.

### Nanoparticle Tracking Analysis

A Nanosight NS300 (Malvern Panalytical, Malvern, UK), equipped with a 532-nm green laser and NS300 FCTP Gasket (Cat no. NTA4137), was utilized for characterizing size and number of circulating exosomes. All samples were analyzed at a camera level of 12 and a detection threshold of 3. Video capture settings were set to record three videos for one min each [47]. Data was analyzed using NTA software v3.2. SC-EV samples were prepared with a 1:140 dilution in a total volume of 1ml of sterile, filtered PBS and injected using a 1 mL sterile BD Plastipak syringe (Becton Dickinson S.A., Madrid, Spain).

### SC-EV isolation, Labeling, and Uptake

EVs were isolated from SC culture media using ExoQuick TC (System Bioscience, Palo Alto, USA), following the manufacturer’s instructions and resuspended in 1X PBS for long term storage at −80°C. Isolated SC-EVs were labeled using 10µM carboxyfluorescein succinimidyl ester (CFSE) dye in PBS for 2 h at 37°C. Following the incubation period, ExoQuick TC was used via manufacturer’s instructions to re-pellet and isolate the labeled SC-EVs. The labeled SC-EVs were then resuspended in 100μl PBS and 50 µg labeled SC-EVs were added to the media of C2C12 in an eight chamber cover glass slide and incubated at 37°C for 24 h. Following the 24 h incubation period, the myotubes were then washed with PHEM buffer (36.28 g PIPES, 11.92 g HEPES, 7.6 g EGTA, 1.97 g MgSO_4 •_ 7H_2_O) to remove any labeled protein that was not incorporated into the myotubes and fixed in 4% paraformaldehyde (PFA) for 10 min at room temperature. Imaging of the SC-EV protein uptake was then performed on the Leica LSM-880 confocal microscope system using the 488 nm laser line and 63X objective lens (Zeiss, Oberkochen, Germany) to obtain a representative image of labeled SC-EV protein uptake in the myotubes.

To quantify the uptake of SC-EVs, C2C12 were cultured in a 96 well micro-clear plate (Greiner Bio-One, Monroe, USA). Separately, EVs were isolated from SC culture media and labeled via incubation with 10µM CFSE at 37°C for 2 h. Excess dye was removed by re-isolating the labeled-EVs using ExoQuick TC. 1.00×10^9^ labeled EVs were then incubated in each well of the 96 well plate for 5 min, 30 min, 1 h, 2 h, 6 h, 24 h and 48 h. Following the incubation period, the myotubes were washed with PHEM buffer to remove any labeled protein that was not incorporated into the myotubes and fixed in 4% PFA for 10 min at room temperature. The average spot intensity measurement on the Cellinsight CX7 (Thermo Fisher Scientific, Waltham, USA) was used to analyze the uptake of CFSE fluorescently labeled SC-EV protein delivered to C2C12 myotubes. The 488 nm laser line and 20X objective lens on the Leica LSM-880 confocal microscope system were then used to obtain representative images of CFSE-labeled SC-EV protein in the myotubes at each timepoint.

### SC-EV protein colocalization with mitochondria

CFSE-labeled SC-EVs were incubated in the differentiation media of C2C12 myotubes for 24 h, as previously described. Following the 24 h incubation period, the media was changed to media containing MitoTracker (Invitrogen, Carlsband, USA) at a final concentration of 300 nM for 45 min in a 5% CO_2_ cell culture incubator. The media was then removed and the myotubes were washed and fixed in 4% PFA solution. The myotubes were then imaged on the Leica LSM-880 confocal microscope system using the 488 nm and 561 nm laser lines and 40x objective lens to obtain a representative image of SC-EV protein and mitochondrial colocalization.

### Live Cell Metabolic Assay

C2C12 myotubes were cultured on eight-well Seahorse culture plates (Agilent, Santa Clara, USA). 1.0 × 10^4^ cells were plated into each well. The wells on each plate were then divided into four different groups: control (no treatment), SC-EV treatment (3.12×10^8^ SC-EV/well), treatment with 50 µM H_2_O_2_, or H_2_O_2_ with subsequent SC-EV treatment (H_2_O_2_ + SC-EV). Each treatment was conducted for 24 h. The H_2_O_2_ treatments were administered first by diluting the H_2_O_2_ in DM and performing a media change from regular DM to the H_2_O_2_-treated DM. Meanwhile, the untreated control and SC-EV treatment groups remained in regular DM. Following 24 h of H_2_O_2_ treatment, all groups received a wash with fresh DM as well as a media change with either regular DM or SC-EV treatment DM. The SC-EV and H_2_O_2_ + SC-EV groups received SC-EV treatment DM containing 3.12×10^8^ SC-EVs/well, whereas the untreated control and H_2_O_2_ groups received fresh regular DM containing no SC-EVs. All conditions were performed in 3 independent experiments in duplicate (3 technical replicants). Following the treatment period, the media was removed, the cells were washed with XFp Assay media and then replaced with fresh XFp Assay media, composed of 7.4 pH XF DMEM (Agilent, Santa Clara, USA), supplemented with 1 mM pyruvate, 2 mM glutamine, and 10 mM glucose. The cells were incubated in the assay media for 45 mins in a CO_2_-free 37°C incubator, and then analyzed on the Agilent Seahorse platform following the manufacturer’s mitochondrial stress test protocol. Briefly, the compounds in the kit were reconstituted to the manufacturer’s specified concentrations: 1.5 µM Oligomycin, 1.0 µM FCCP, and 0.5 µM rotenone A. Pre-specified amounts of these compounds were then loaded into the ports of the extracellular flux cartridge provided in the kit. The machine then exposed the myotubes to each compound in a sequential order while measuring various parameters of mitochondrial function.

### Statistical Analyses

When results from two groups were compared, a *t*-test was used to test for significance. When results from more than two groups were compared, a one-way ANOVA was used to determine overall significance. When appropriate, a post hoc Tukey honestly significant difference (HSD) test was performed. Differences in results were considered significant when P ≤ 0.05. For each outcome, at least 3 samples per treatment group, acquired from ≥ 3 independent experiments, were quantified and analyzed.

## Acknowledgements

Supported by NIH R01 NS102157, NIH P20 GM113125, NIH P20 GM103446, NIH R03 HD094594 to MBH

---

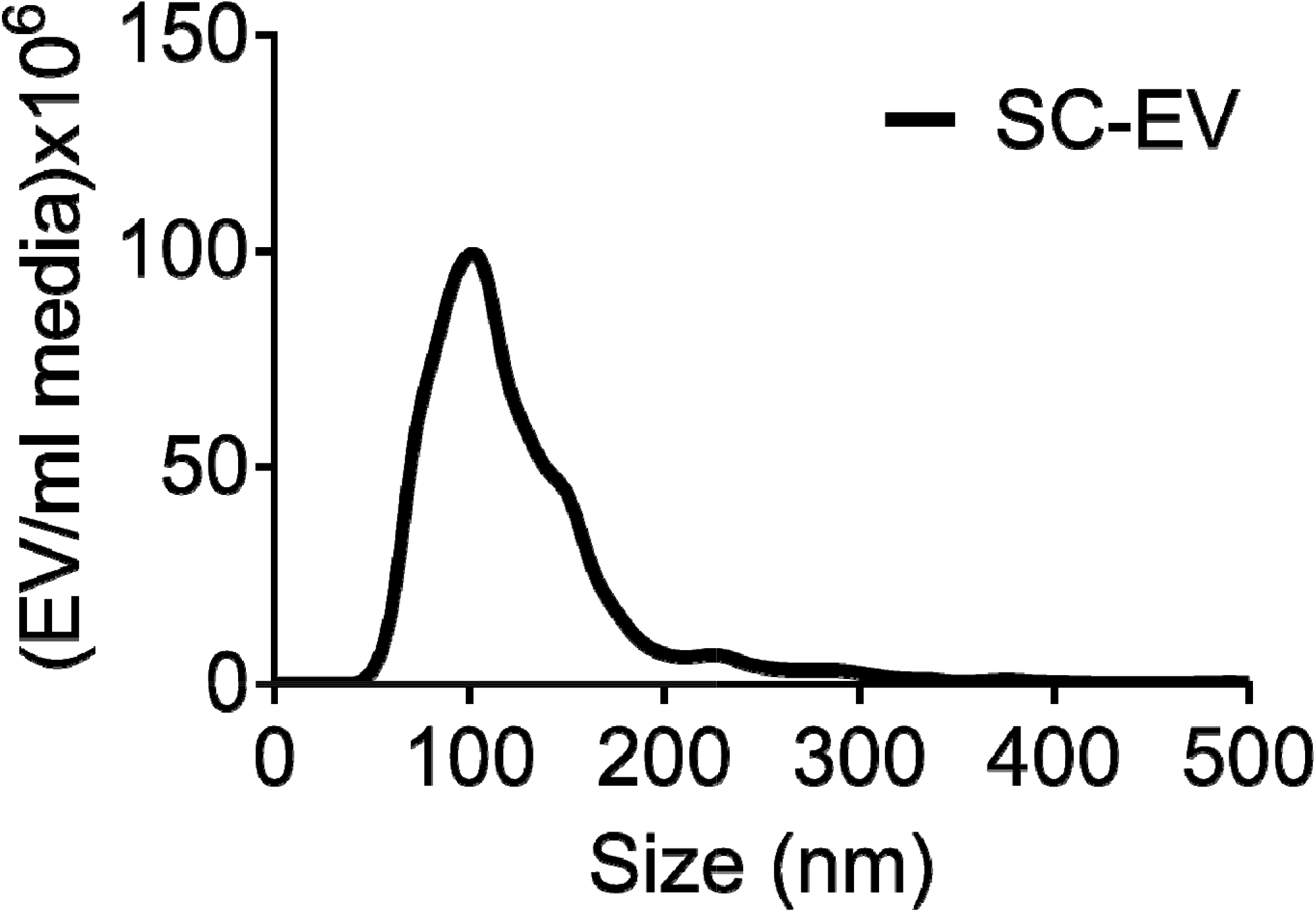

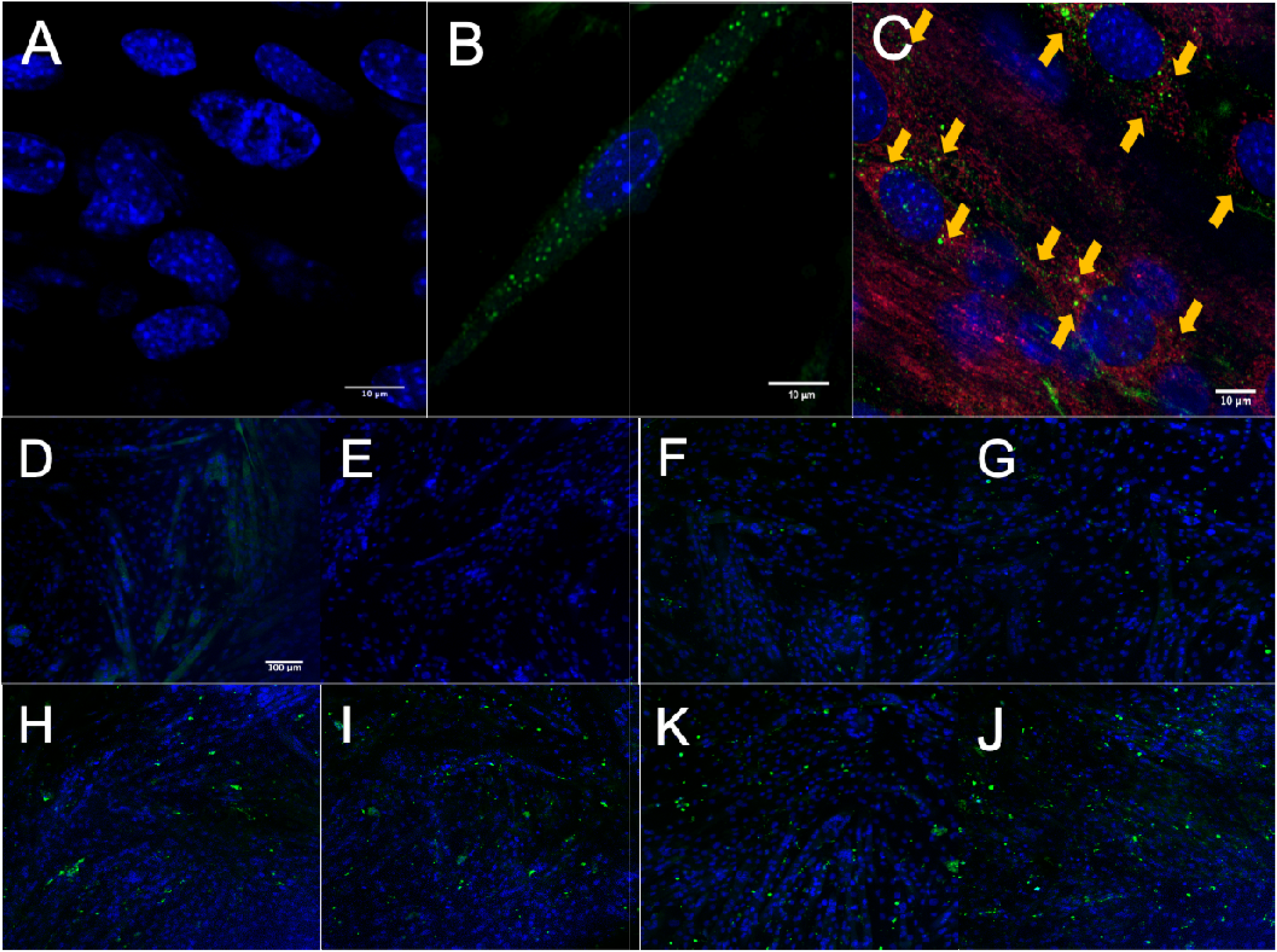

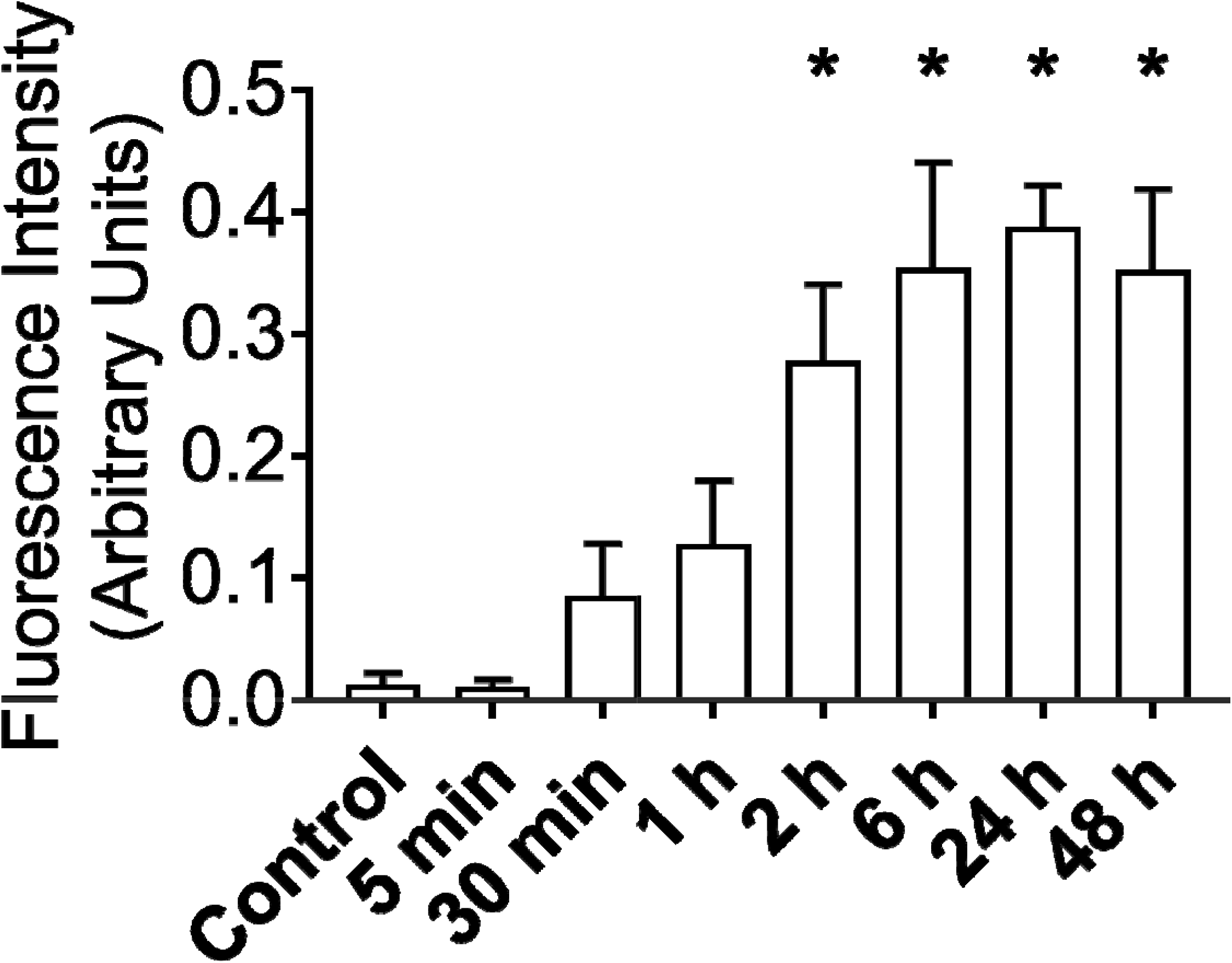

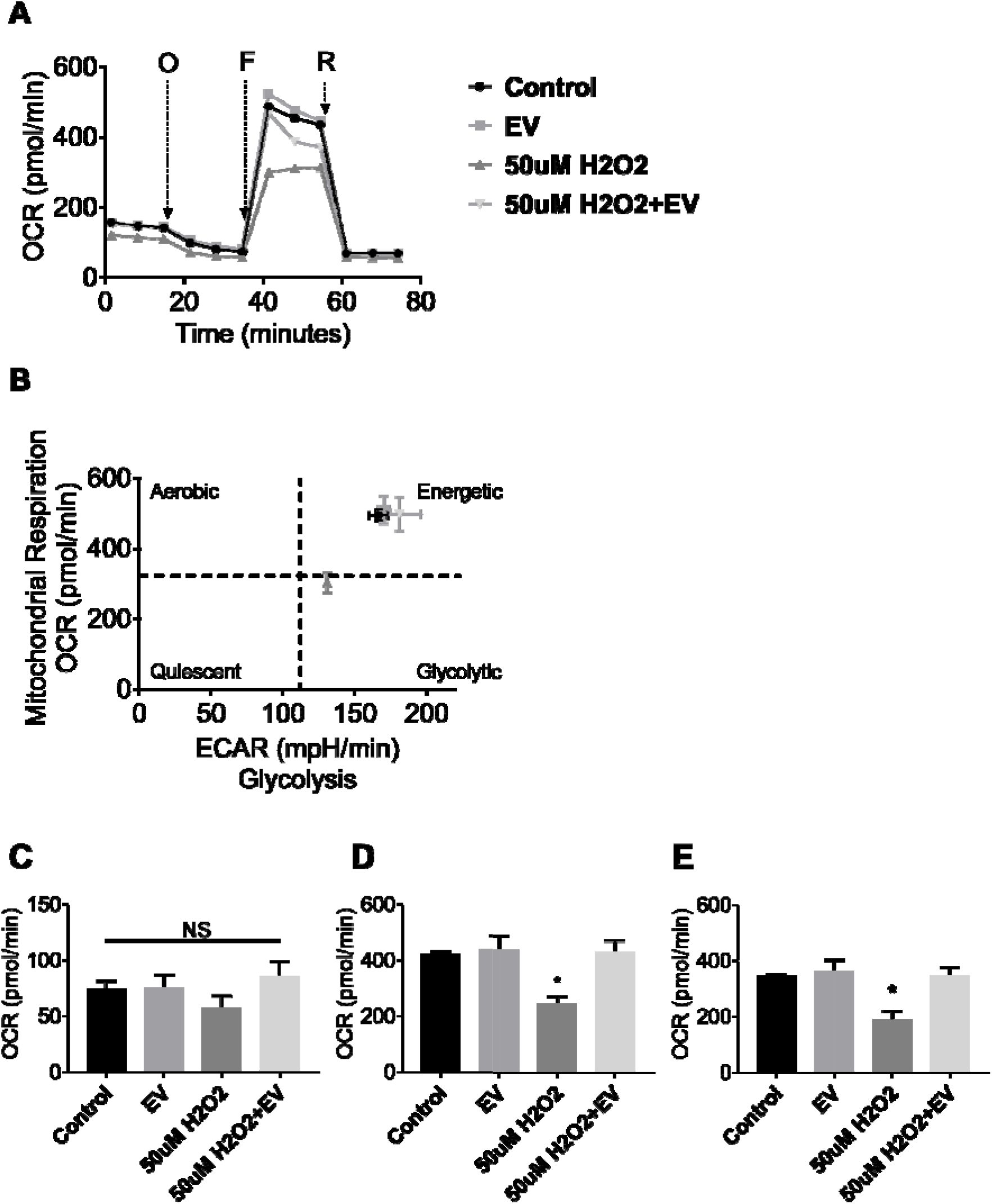

